# Maize *rough endosperm6* is a predicted RNA helicase required for miRNA processing and endosperm cell patterning

**DOI:** 10.1101/2024.12.13.628378

**Authors:** Tianxiao Yang, Masaharu Suzuki, L. Curtis Hannah, A. Mark Settles

## Abstract

Maize *rough endosperm* mutants have defective kernels with a rough, etched, or pitted endosperm surface. Molecular genetic analysis of this mutant class has identified multiple RNA processing proteins critical to endosperm development. Here, we report that *rough endosperm6* (*rgh6*) encodes a predicted DEAD-box RNA helicase required for miRNA processing. Mutant *rgh6* kernels reduce grain fill and increase relative transcript levels of markers specific to epidermal endosperm cell types. B-A translocation crosses revealed that *rgh6* mutant endosperm inhibits normal embryo development. We mapped the *rgh6* locus to a three gene interval and confirmed it encodes a predicted DEAD-box RNA helicase with two independent transposon-tagged alleles. Transient expression of a RGH6-green fluorescent protein (GFP) fusion is localized to nucleolus and nuclear speckles consistent with a function in nuclear RNA processing. Mutant *rgh6* endosperms have increased precursor miRNA and decreased mature miRNA levels indicating that *rgh6* impacts miRNA processing. Our study demonstrates that precursor miRNA processing and miRNA target regulation are required for normal endosperm development.

## Introduction

The endosperm is a vitally important, multicellular tissue within angiosperm seeds and is required for normal embryo development. In maize, endosperm development is initiated when the diploid central cell is fertilized by a haploid sperm cell. Early endosperm is a syncytium generated from free nuclear division without cell wall formation. Syncytial endosperm transitions to cellular endosperm via alveolation (Sabelli and Larkins, 2009). As the endosperm proliferates, cells differentiate along the radial axis to form internal starchy endosperm (Woo et al. 2001) and multiple epidermal cell types. Endosperm epidermal cells differentiate based on the micropylar-chalazal axis to form the basal endosperm transfer layer (BETL) (Hueros et al. 1999; Muñiz et al. 2006), the aleurone layer (Gómez et al. 2009), the embryo-surrounding region (ESR) (Opsahl-Ferstad et al. 1997), and the endosperm adjacent to scutellum (EAS) (Doll et al. 2020). Hormones, sugars, metabolites, peptide-receptor signaling, and transcription factors affect endosperm cell developmental fates (Doll et al. 2017).

Plant microRNAs (miRNAs) are important for gene regulation through silencing of target mRNAs (Rogers and Chen, 2013; Yu et al. 2017; Li and Yu, 2021; Xu and Chen, 2023). Two studies demonstrate a role for miRNAs in maize endosperm development. First, miR169o positively regulates mature kernel weight and volume (Zhang et al. 2022). The NF-YA13 transcription factor is a direct target of miR169o and affects the size and number of starchy endosperm cells. NF-YA13 binds to the promoter and controls the expression of an auxin synthesis gene, YUC1, which in turn modulates auxin homeostasis and cell division. Similarly, miR159 promotes endosperm cell number and mature kernel size by degrading mRNA encoding the MYB74 and MYB138 transcription factors (Wang et al. 2023).

MicroRNA genes (*MIRs*) are transcribed by RNA polymerase II (Pol II) into primary miRNAs (pri-miRNAs). The pri-miRNAs are first processed into precursor miRNAs (pre-miRNAs), which are then processed into the miRNA/miRNA* duplexes. The processing steps are catalyzed by the dicing complex, which includes the ribonuclease III protein DICER-LIKE 1 (DCL1) to cleave miRNA precursors as well as the double-stranded RNA-binding protein HYPONASTIC LEAVES 1 (HYL1) and the zinc-finger protein SERRATE (SE) that ensure cleavage accuracy (Dong et al. 2008). DCL, HYL1, and SE are co-localized into nuclear loci called Dicing bodies (D-bodies) (Fang et al. 2007; Song et al. 2007). Weak alleles of the maize *dcl1* gene cause the *fuzzy tassel* (*ftz*) plant developmental phenotypes, while strong alleles give rise to lethal embryo and endosperm defects showing an essential role for miRNA processing in maize seed development (Thompson et al. 2014; Xie et al. 2023).

RNA helicases are ubiquitous and essential proteins in nearly all aspects of RNA metabolism, including transcription, RNA transport, translation, RNA decay, and RNA processing. RNA helicases are enzymes that bind or remodel RNA, ribonucleoprotein (RNP) complexes, or both using ATP (Cordin et al. 2006; Fairman-Williams et al. 2010). Together with DNA helicases that are involved in replication, recombination and repair, all helicases can be classified into six superfamilies, from SF1 to SF6, based on sequence and structural features. The SF1 and SF2 families are present in eukaryotic genomes, whereas SF3 to SF6 are found only in bacteria and viruses (Jankowsky, 2011). The eukaryotic RNA helicases contain a helicase core of two tandem recombinase A (RecA)-like domains. The helicase core is flanked by variable auxiliary N- and C-terminal extensions. Within the helicase core, at least 12 characteristic sequence motifs are located at conserved positions. The helicase motif II, which has the Asp-Glu-Ala-Asp (D-E-A-D) amino acid sequence, has inspired the name of DEAD-box for this family. Helicase motifs correspond to the primary function of ATP binding and hydrolysis, RNA binding, and coupling ATP and RNA binding (Linder and Jankowsky, 2011).

DEAD-box RNA helicases are also required for pri-miRNA processing. DEAD-box RNA helicase 6 (RH6), RH8, and RH12 are D-body components (Li et al. 2021), while RH27 is required for normal transcript levels of a subset of pri-miRNAs (Hou et al. 2021). The DEAD-box RNA helicase, RCF1, is required for pri-miRNA processing and splicing, and as well as pre-mRNA splicing. RCF1 enhances D-body formation and facilitates the interaction of pri-miRNAs and HYL1 (Xu et al. 2023).

Mutations in genes required for different miRNA processing steps typically result in reduced levels of mature miRNA. For example, mutants in *se* (Li et al. 2020), *rh27* (Hou et al. 2021), or *rcf1* (Xu et al. 2023) reduce miRNA levels. However, *rh27* mutants show decreased levels of pri-miRNA, suggesting a role in primary miRNA transcript accumulation, while *se* and *rcf1* have increased levels of pri-miRNA indicating roles downstream of pri-mRNA production (Li et al. 2020; Hou et al. 2021; Xu et al. 2023).

Wang et al. (2024) recently reported that maize *defective kernel51* (*dek51*) is a nucleolus-localized DEAD-box RNA helicase. Mutation in *dek51* shows severe embryogenesis defects with impaired developmental transition, and *dek51* embryos have aberrant 3’ end processing intermediates of both 18S and 5.8S pre-ribosomal RNA (pre-rRNA), although mature rRNA levels are unaffected. A role in pre-rRNA processing was proposed due to protein-protein interaction assays suggesting DEK51 can interact with putative factors of the nuclear RNA exosome and putative pre-rRNA endonucleases.

Here, we show that maize *rough endosperm6* (*rgh6*) encodes the identical DEAD-box RNA helicase as maize *defective kernel51* (*dek51*). Like *dek51*, we observed no impact on mature rRNA levels in *rgh6* endosperm. However, we found that *rgh6* affects processing of mature miRNA. Mutation in *rgh6* causes reduced endosperm size with increased transcript levels of epidermal cell type markers. Genetic mosaics and *rgh6* mutant embryo phenotypes suggest the normal allele of *Rgh6* is required in the endosperm to support embryo development. By focusing on *rgh6* endosperm phenotypes, this study revealed developmental roles for maize miRNAs in endosperm epidermal cell differentiation.

## Results

### Inheritance of *rgh6*

The *rgh6* mutation was isolated from the UniformMu population, which is a Robertson’s *Mutator* (*Mu*) transposon-tagging population in the color converted W22 inbred (McCarty et al. 2005). Self-pollination of heterozygous *rgh6* plants produced normal and *rgh* kernel phenotypes at a ratio of 3 to 1, suggesting that *rgh6* acts as a recessive mutation (Figure 1A). When heterozygous *rgh6* plants were crossed as the female or male parent with the W22 inbred, all kernels are visually normal indicating that *rgh6* is unlikely to have a gametophyte effect on seed development (Figure 1B and 1C).

**Figure 1.**
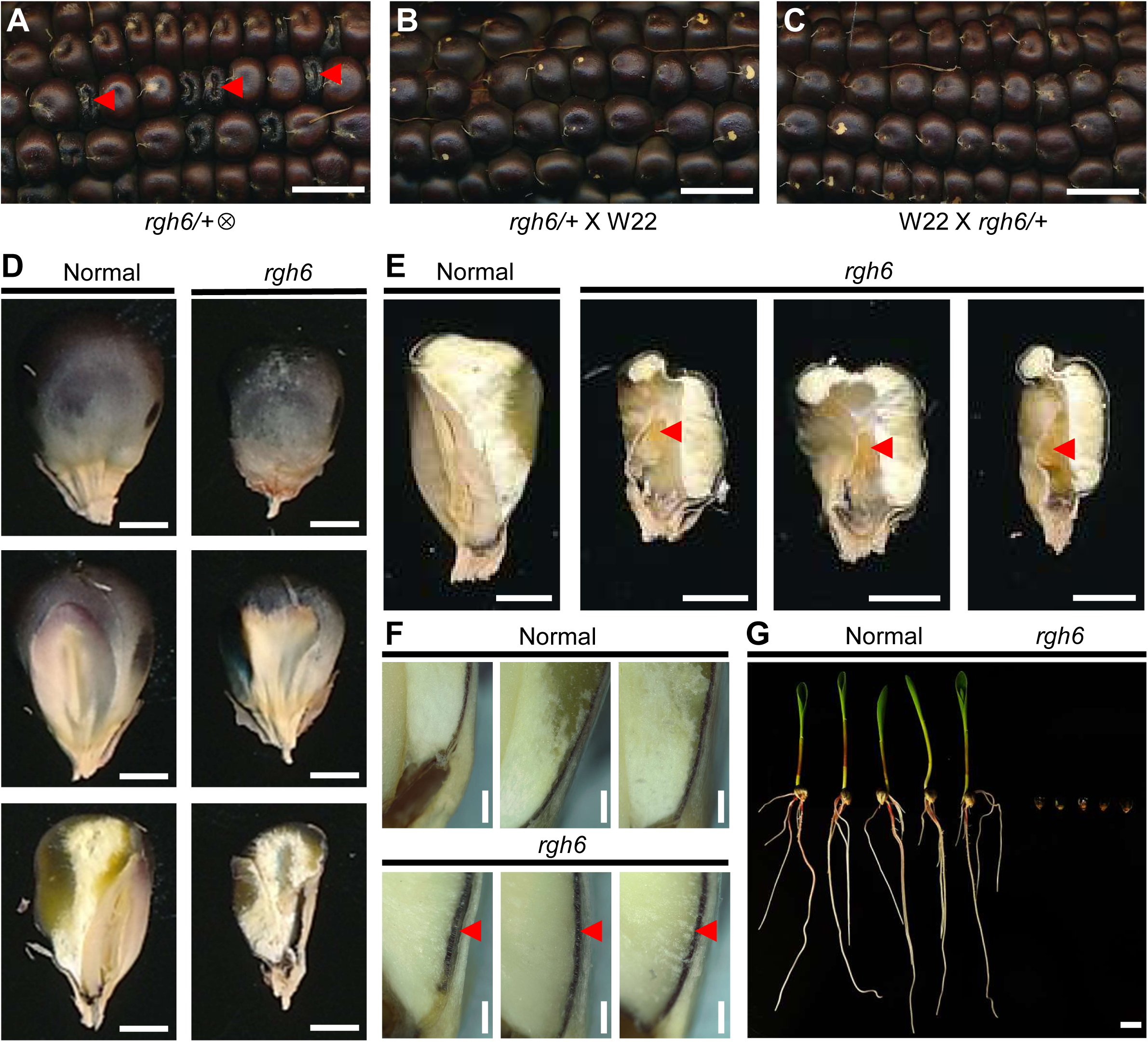
Kernel phenotypes of *rgh6* mutant. (A) Self-pollinated ear segregating for *rgh6-umu1* in the UniformMu background. Arrowheads indicate *rgh* kernels. Scale bar is 1 cm. (B-C) Reciprocal crosses of *rgh6-umu1* heterozygotes with color converted W22. Second ears were self-pollinated to identify *rgh6-umu1* heterozygotes. Scale bars are 1 cm. (D) Top, middle and bottom panels are the abgerminal view, germinal view and sagittal section of normal and *rgh6-umu1* sibling kernels. Scale bars are 0.25 cm. (E) Sagittal sections of *rgh6-umu1* and normal siblings. Arrowheads indicate embryo position. Scale bars are 0.25 cm. (F) Aleurone (arrowheads) of *rgh6* mutants and normal siblings. Scale bars are 0.5 mm. (G) Germination experiments for *rgh6* mutants and normal siblings. Scale bar is 1 cm.

### Kernel phenotype of *rgh6*

Homozygous *rgh6* mutants are reduced grain-fill, defective kernels (Figure 1D). Compared to normal kernels, *rgh6* endosperm has a smaller fraction of vitreous endosperm and a larger air space at the basal region (Figure 1D). Mutant *rgh6* embryos generally arrest at a globular developmental stage consistent with a reduced oil content at kernel maturity (Figure 1D). Defective *rgh6* embryos develop in a more apical position than typical for other defective embryo mutants (Figure 1E). For example, two alleles of *defective kernel5* (*dek5*), *dek5-PS25* and *dek5-MS33,* have mutant embryos at the basal position of mature kernels (Zhang et al. 2019). Although the *rgh6* aleurone has a normal single cell layer, the mutant aleurone cells are elongated in the radial axis compared to normal siblings (Figure 1F). The *rgh6* kernels do not germinate (Figure 1G).

### *rgh6* endosperm negatively impacts normal embryo development

Maize B-A translocation stocks produce genetically non-concordant endosperm and embryo tissues when crossed as males (Roman, 1947). These stocks contain translocations of normal, A chromosome arms with a supernumerary B centromere that undergoes non-disjunction in the second pollen mitosis to produce aneuploid sperm cells, which are either hyperploid or lacking the translocated A chromosome. Double fertilization of a recessive kernel mutant produces genetic chimeras with the defective locus uncovered in either endosperm or embryo. Endosperm-embryo interactions can be inferred from hand sections of non-concordant kernels. Mutants are considered to have autonomous endosperm and embryo phenotypes when two classes of non-concordant kernels are observed, either defective in endosperm or in embryo. By contrast, uncovered mutants are considered to have non-autonomous interactions when the non-concordant kernels are defective in both endosperm and embryo (Fouquet et al. 2011).

We generated genetically non-concordant kernels by crossing *rgh6* heterozygous plants as female parents with a hyperploid TB-5Sc B-A translocation stock (Birchler and Alfenito, 1993). Two classes of uncovered kernels were found. One class had a defective embryo with normal endosperm, and the other class showed defects in both the endosperm and embryo (Figure 2A). As a non-autonomous control, the TB-5Sc stock was crossed onto *rgh*-744* (Fouquet et al. 2011). The *rgh*\*-744 crosses also showed two classes of uncovered kernels in which a genetically normal embryo is not able to develop with a mutant endosperm (Figure 2B).

**Figure 2.**
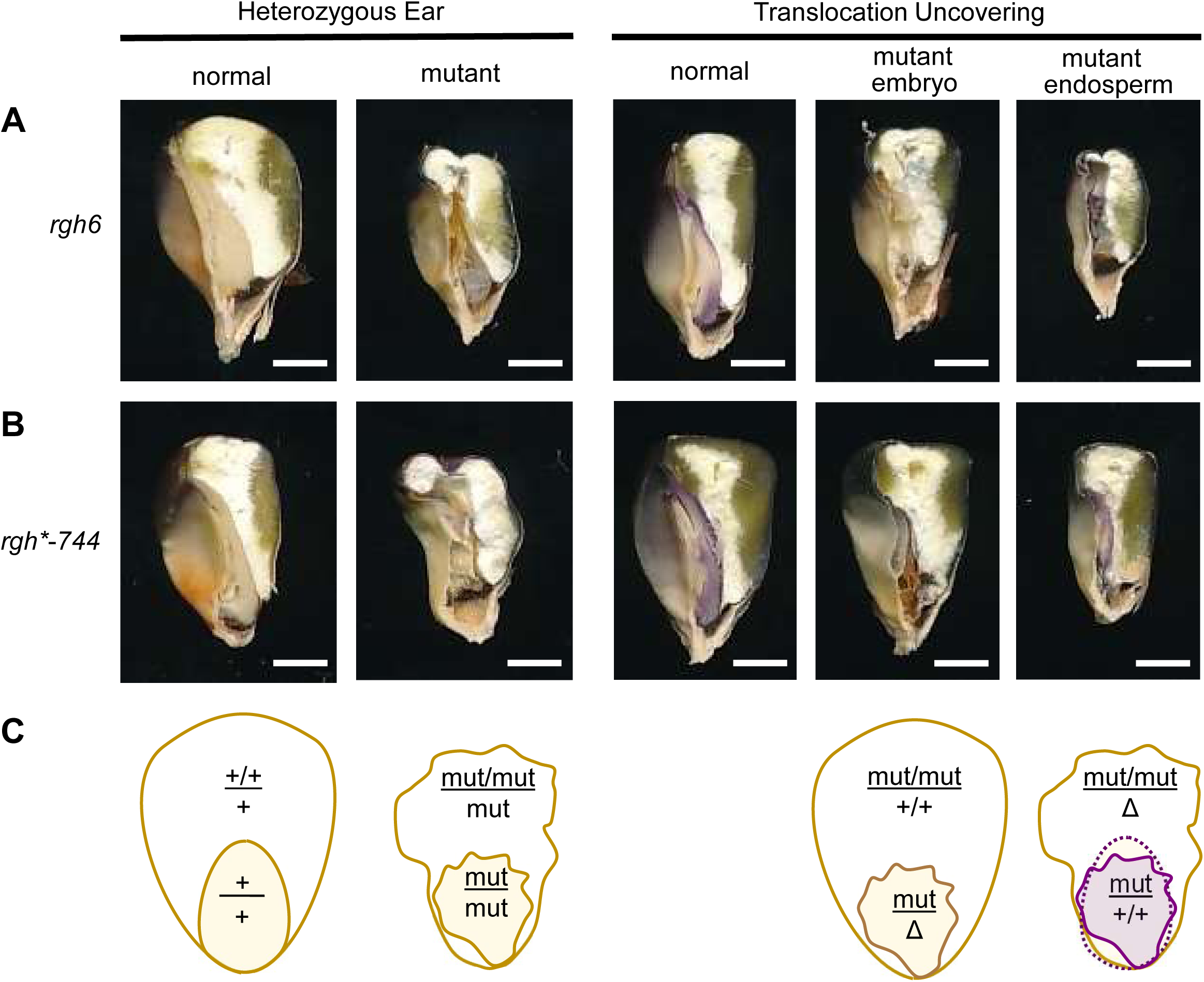
*rgh6* mutant endosperm inhibits normal embryo development. (A-B) Sagittal sections of mature kernels from a self-pollinated heterozygous ear and TB-5Sc translocation crosses of (A) *rgh6* and (B) *rgh*-744* heterozygous plants. Scale bars are 0.25 cm. (C) Schematic of inferred genotypes based on developmental and anthocyanin phenotypes of the kernel images. Aneuploidy lacking the of the short arm of chromosome 5 is represented by Δ.

We used anthocyanin color markers to confirm transmission of the TB-5Sc translocation into embryos of uncovered *rgh6* kernels. TB-5Sc is marked with a dominant *A2* allele that expresses anthocyanins, and the stock carries an *R1-scm2* allele. Combined these alleles give purple color to embryos when the TB-5Sc sperm fertilizes the egg cell (Figure 2C). In *rgh6* and *rgh*-744*, uncovered kernels with mutant endosperm and defective embryos were purple, reporting that these seeds were fertilized with the translocation stock (Figure 2A and 2B). These results indicate that an embryo with a functional *Rgh6* allele fails to develop when the endosperm is mutant for *rgh6*.

### *Rgh6* encodes a predicted DEAD-box RNA helicase

We generated an F_2_ mapping population from heterozygous *rgh6* crossed with the B73 inbred and mapped with distributed simple sequence repeat (SSR) markers (Martin et al. 2010). SSR markers linked to *rgh6* are expected to show segregation distortion for the W22 allele among F_2_ mutant kernels (Supplementary Figure S1A). The hybrid F_2_ ears segregated for *rgh6* kernels at a 3 to 1 ratio indicating a single-locus recessive phenotype (Supplementary Figure S1B and S1C). Consistent with TB-5Sc uncovering crosses, a genome scan of SSR markers localized the *rgh6* reference allele to the short arm of chromosome 5 (Figure 3A). Fine mapping narrowed the *rgh6* reference allele to a 60 kbp genomic interval with three gene models in the B73 RefGen_v3 genome assembly (Figure 3B). One gene model, GRMZM2G350428, encodes a CLAVATA-LIKE protein but is not expressed in the endosperm tissue based on an RNA-seq gene atlas and was eliminated from further consideration (Stelpflug et al. 2016). We amplified genomic DNA for the additional two gene models from homozygous *rgh6* mutants and found a *Mu* transposon insertion in the predicted DEAD-box RNA helicase gene, GRMZM5G897976, that co-segregated with the *rgh* kernel phenotype (Figure 3C and 3D). This *rgh6* reference allele was named *rgh6-umu1*.

**Figure 3.**
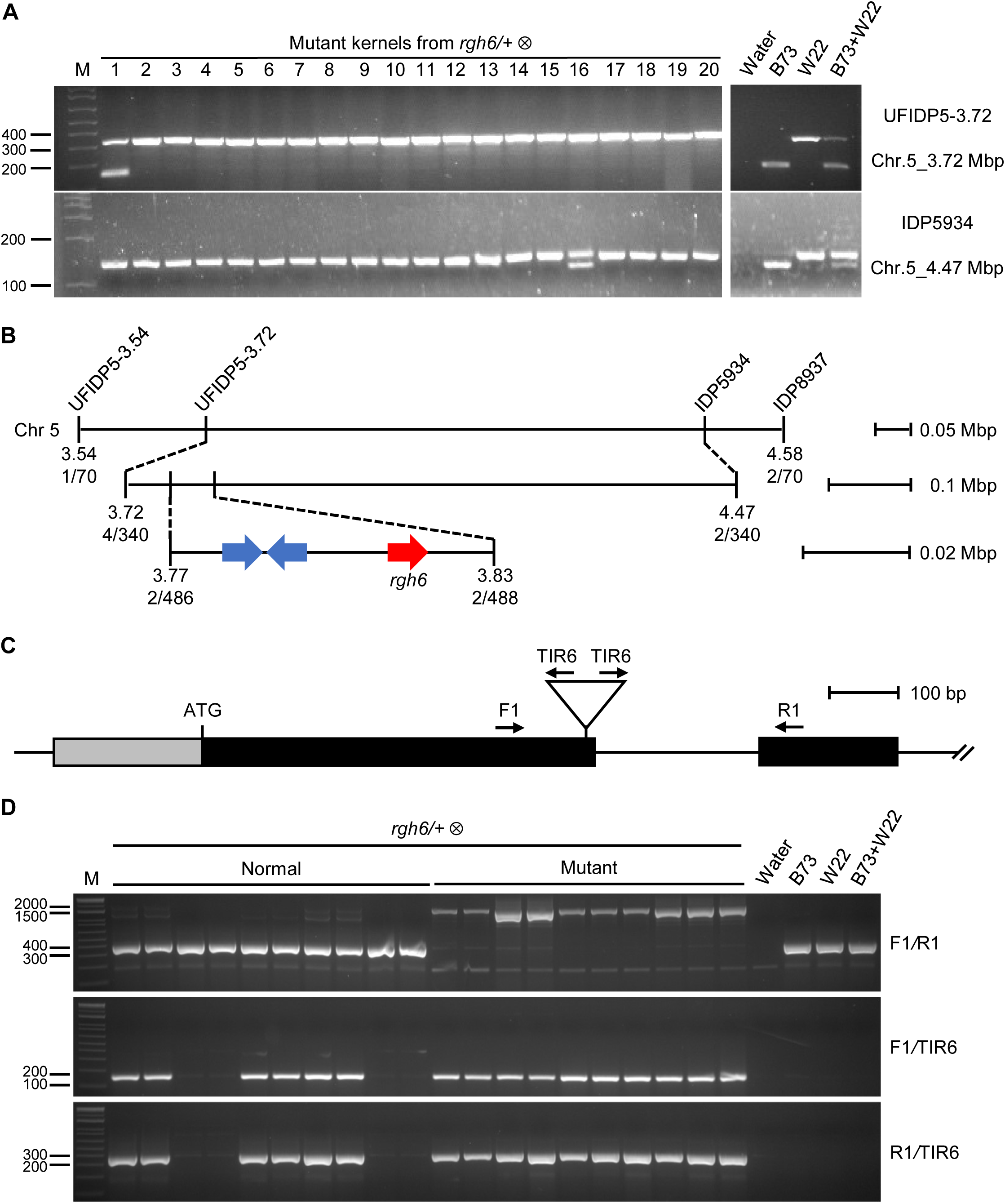
Positional cloning of *rgh6* locus. (A) Flanking IDP markers showing recombinants from individual mutant kernels in the F_2_ mapping population. (B) Schematic of fine mapping results. Molecular markers are indicated above. B73 RefGen_v3 genome assembly physical positions are in megabases (Mb). Genetic recombination distances are in centimorgans (cM). Gene models are denoted by arrows with the *rgh6* gene in red. (C) Schematic of partial *rgh6* locus and primers used for PCR analysis. F1 and R1 are the *rgh6* locus specific primers, and TIR6 is the *Mu* transposon primer. (D) Co-segregation analysis of *rgh6-umu1* with the *rgh6* phenotype in a segregating self-pollination.

A second allele, *rgh6-umu2*, was available in a UniformMu stock accession, UFMu-11204, segregating for the insertion site, mu1095038::*Mu* (Figure 4A). *Mu* insertion genomic flanking PCR products were amplified from *rgh6-umu1* and *rgh6-umu2*, and the product sequences revealed independent *Mu* insertions at the identical position in exon 1. The insertions for each allele have polymorphic terminal inverted repeat (TIR) sequences between *rgh6-umu1* and *rgh6-umu2*, indicating insertion of two different *Mu* elements (Figure 4B). Self-pollination of *rgh6-umu1* and *rgh6-umu2* heterozygotes segregated for the *rgh* phenotypes. When these heterozygotes were crossed as male parents onto the W22 inbred, no *rgh* kernels were observed, but reciprocal crosses between these heterozygotes failed to complement the *rgh6* phenotype indicating the *rgh6* locus is encoded by GRMZM5G897976 (Figure 4C).

**Figure 4.**
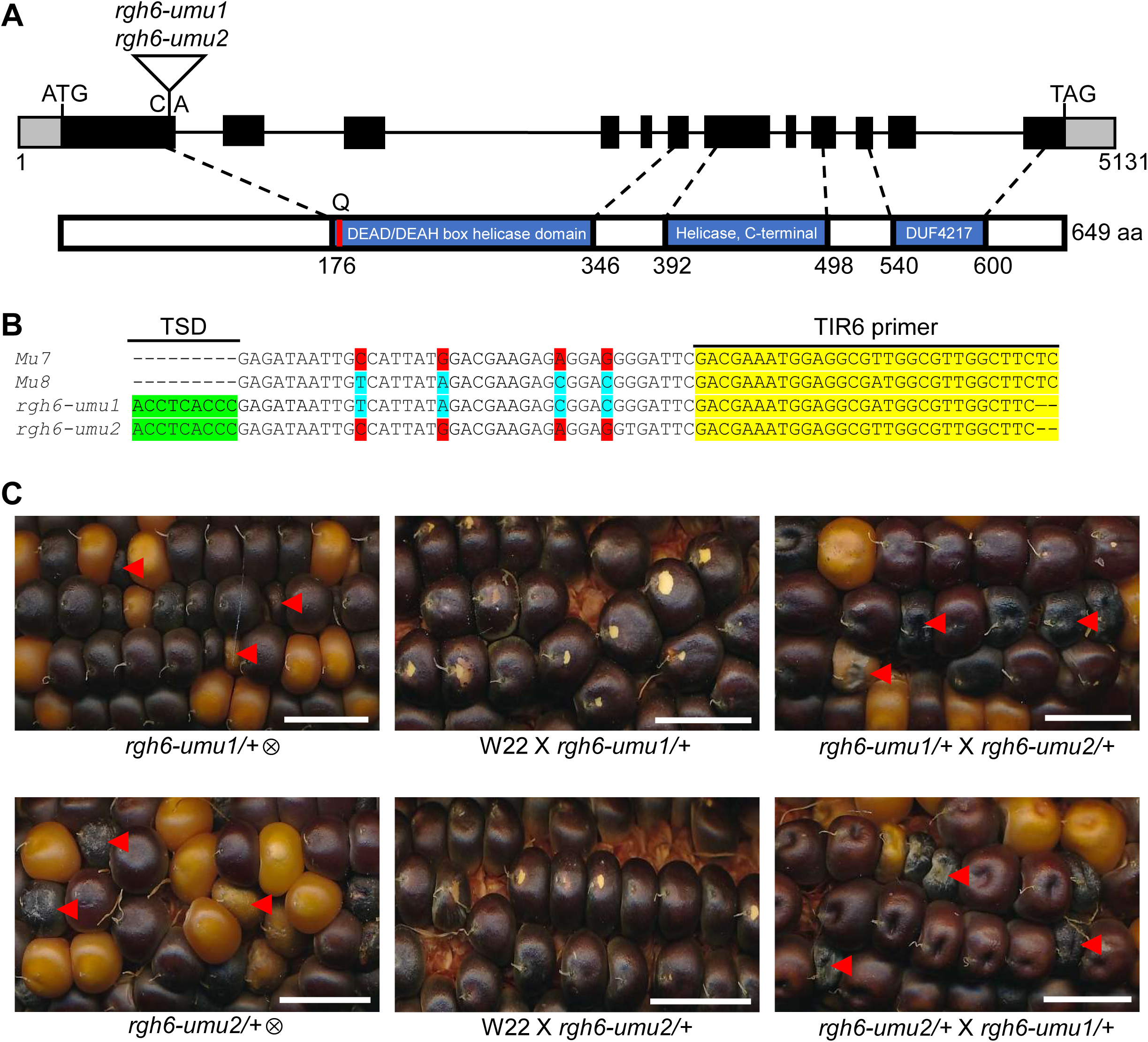
Mutant alleles of *rgh6* locus. (A) Schematic of the *rgh6* gene and protein. Black boxes are coding sequence exons, gray boxes are untranslated exon, and black lines are introns. White triangle in the gene indicate the *Mu8* and *Mu7* insertions at the same genomic position in *rgh6-umu1* and *rgh6-umu2*, respectively. The red line in the protein indicates a Q motif in the DEAD/DEAH box helicase domain. (B) Sequence alignments of *Mu7* and *Mu8* reference sequences with *Mu* flanking sequences from *rgh6-umu1* and *rgh6-umu2* amplified with the Rgh6-F1 and TIR6 primers. TSD is the *Mu* target site duplication. (C) Genetic complementation test with control crosses. Self-pollinations (⊗) show *rgh* kernel phenotypes for both alleles. W22 crosses of *rgh6-umu1* and *rgh6-umu2* heterozygotes do not give *rgh* kernel phenotype. Reciprocal crosses between *rgh6-umu1* and *rgh6-umu2* heterozygotes segregate *rgh* kernels. Arrowheads indicate *rgh* kernels. Scale bars are 1 cm.

According to the maize RNA-seq gene atlas, *Rgh6* is expressed in root, leaf, stem, tassel, and seed tissues (Stelpflug et al. 2016). Additional published transcriptome data showed that *Rgh6* transcripts can be detected during whole seed development from 0 to 38 days after pollination (DAP) with peak levels at 8 DAP (Chen et al. 2014), and Yi et al. (2019) showed that *Rgh6* transcripts can be detected in the nucellus and seed tissues from 0 to 144 hours after pollination (HAP) with a peak at 138 HAP.

### RGH6 is localized to nuclear speckles

The RGH6 protein is predicted to localize in the nucleus using WoLF PSORT (Horton et al. 2007), SignalP-6.0 (Teufel et al. 2022) and TargetP-2.0 (Armenteros et al. 2019) algorithms. Based on the cNLS Mapper tool, RGH6 has a monopartite nuclear localization signal (NLS) near the N-terminus (PSKKRKQPVI) and a bipartite nuclear localization signal (NLS) near the C-terminus (KNPPKVNLDLDSSAAKHRKKMRR) of the predicted protein sequence (Kosugi et al. 2008; Kosugi et al. 2009 a; Kosugi et al. 2009 b).

To test the subcellular localization predictions, we transiently expressed a RGH6-green fluorescent protein (GFP) fusion in *Nicotiana benthamiana* leaves. RGH6-GFP is localized to the nucleolus and punctate foci of varying size within the nucleus (Figure 5A). Analysis of 50 cells expressing RGH6-GFP showed that 13 cells had GFP signal only in the nucleolus, while 37 had GFP signal localized to nucleolus and nuclear speckles (Figure 5A). Co-expression of RGH6-GFP with a nucleolus marker, Arabidopsis PRH75 (Lorković et al. 2004), confirmed that RGH6 co-localizes with the nucleolus (Figure 5B).

**Figure 5.**
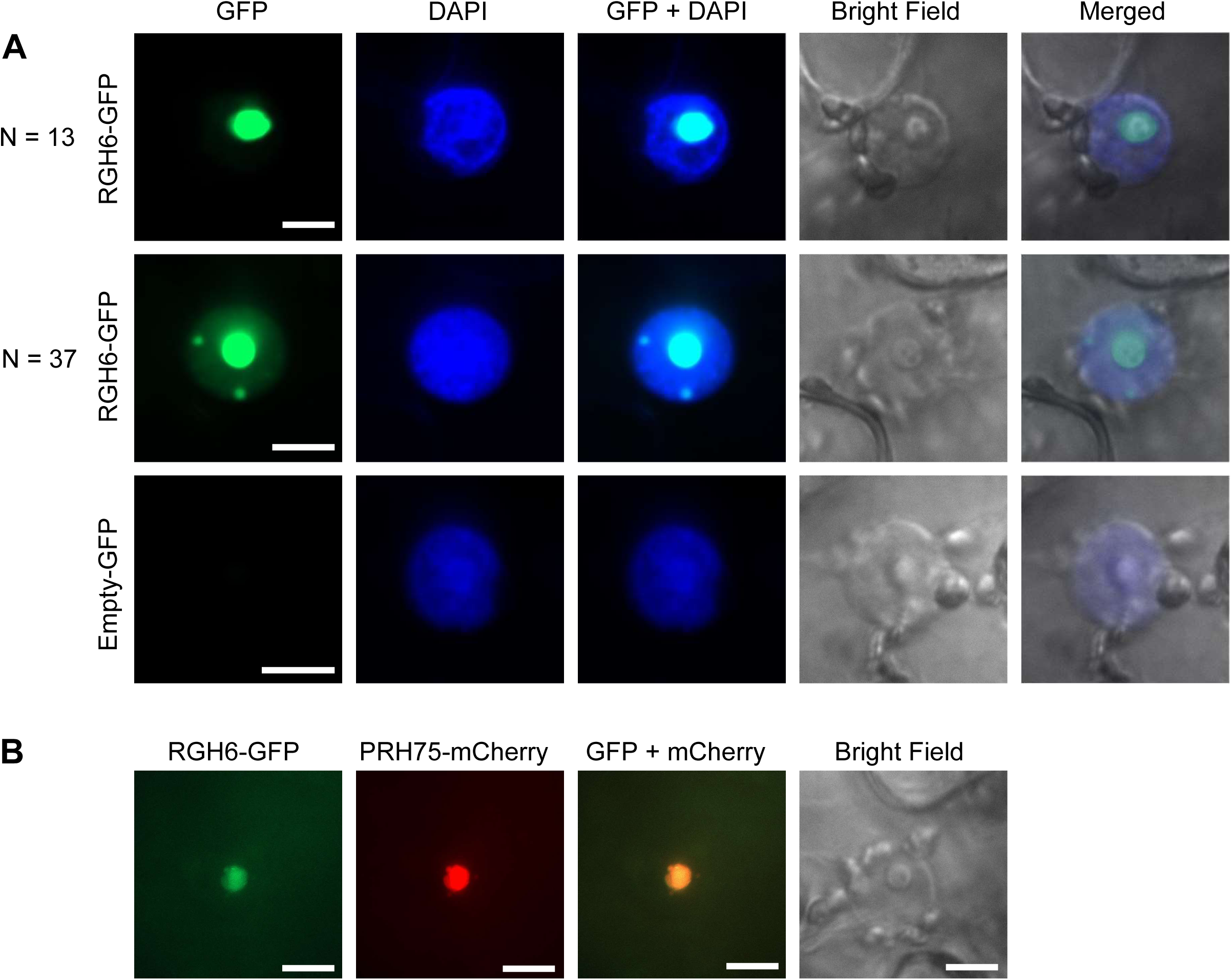
Localization of RGH6-GFP protein. (A) Transient expression of RGH6-GFP fusion and GFP alone in DAPI-stained *N. benthamiana* leaves. N is the number of GFP localization patterns observed in 50 nuclei. (B) Transient co-expression of maize RGH6-GFP and the Arabidopsis PRH75-mCherry nucleolar marker in *N. benthamiana* leaves. Scale bars are 10 µm in all panels.

RNA processing proteins are frequently localized in nuclear speckles and can be localized to the nucleolus in response to cellular stress (Tillemans et al. 2005; Koroleva et al. 2009). A similar localization pattern was observed when the maize minor spliceosome factor, RGH3α, was fused to GFP and expressed in Arabidopsis protoplasts (Fouquet et al. 2011). RH27, a RGH6 homolog in Arabidopsis, is also localized to nucleolus (Hou et al. 2021). Thus, RGH6 subcellular localization is consistent with a function in nuclear RNA processing.

### *rgh6* is unlikely to affect rRNA accumulation and mRNA splicing

To test whether *rgh6* affects rRNA processing, total RNA extracted from endosperm tissue in *rgh6* mutant and normal siblings at 10, 12, and 14 DAP was analyzed by an Agilent 2100 Bioanalyzer. This method showed sharp 25S rRNA and 18S rRNA bands of comparable intensity for the mutant and normal samples (Figure 6A). The relative levels of 25S and 18S were indistinguishable in *rgh6* mutant and normal siblings suggesting that *rgh6* is unlikely to affect rRNA biogenesis (Figure 6B and 6C).

**Figure 6.**
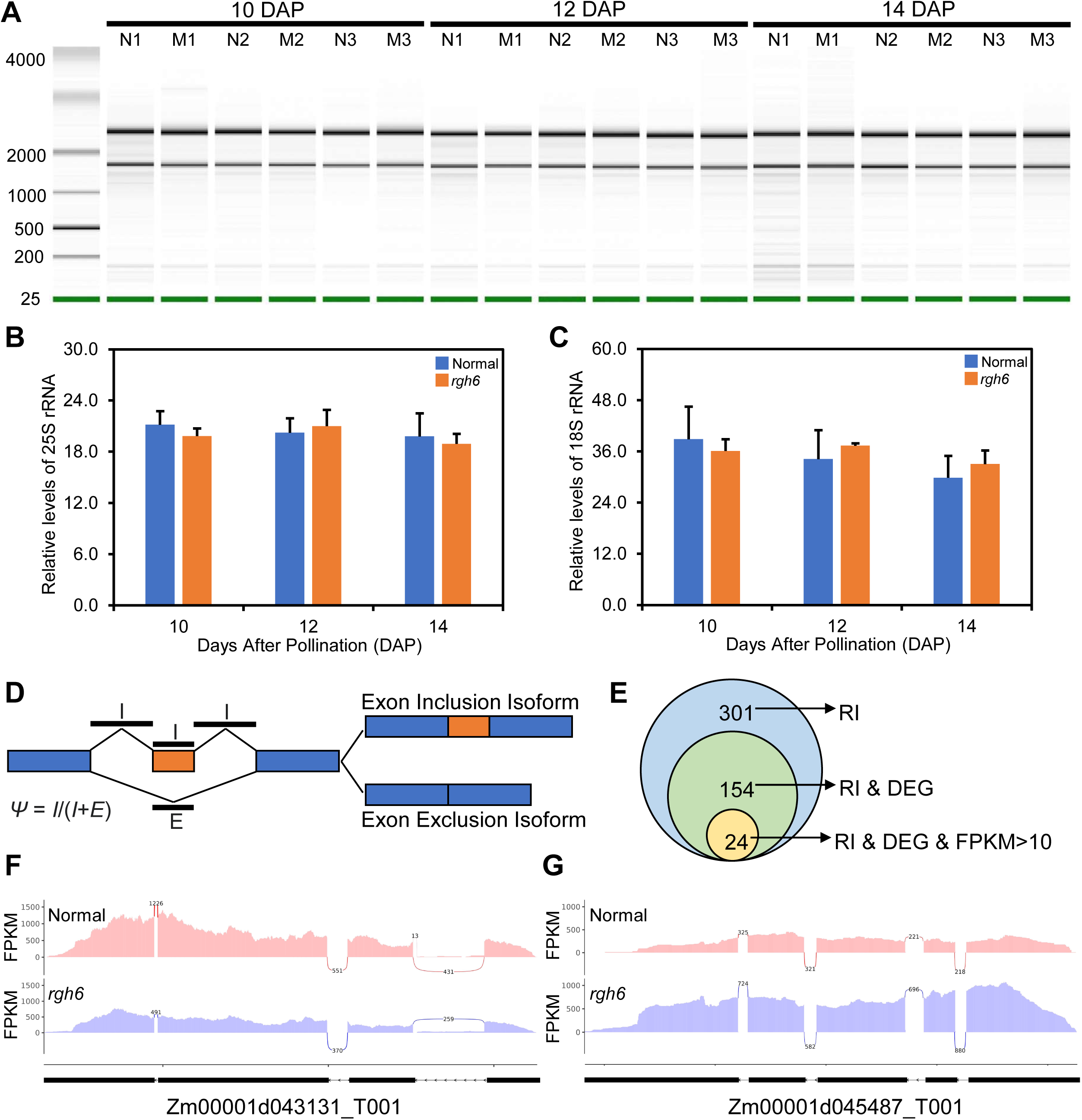
Ribosomal RNA biogenesis and global intron splicing is not affected in *rgh6* endosperm. (A) Agilent 2100 Bioanalyzer results for three biological replicates of normal (N) and *rgh6* (M) sibling endosperm RNA extractions. Green lines indicate a 25 bp internal control. (B) Mean and standard deviation of 25S rRNA levels in *rgh6* mutant and normal sibling endosperm tissues. (C) Mean and standard deviation of 18S rRNA levels from endosperm tissue in *rgh6* mutant and normal siblings. (D) Schematic of exon inclusion level (Ψ) calculation using the fraction of exon inclusion (I) reads relative to the sum of exon inclusion and exclusion (E) reads. The I reads include up- and downstream splice junctions and the alternative exon, while E junction reads span the splice junction from skipping the alternative exon. (E) Venn diagram showing the overlap of significant RI, DEG, and highly expressed, FPKM>10, genes from the mRNA-seq dataset. (F-G) Examples of down-regulated (F) and up-regulated (G) DEGs with significant RI events based on rMATS.

We then completed mRNA-seq from the mutant and normal 10 DAP endosperm tissues. Principal Components Analysis (PCA) of the total gene expression levels showed the RNA samples for the *rgh6* mutant were distinct from normal siblings based on Principal Component 1 (PC1), which explains 53% of the variation in gene expression between samples (Supplementary Figure S2A). A total of 2,270 up-regulated and 1,176 down-regulated genes were differentially expressed with an FPKM >1, fold change >2, and adjusted p-value <0.05 (Supplementary Figure S2B). Gene Ontology (GO) enrichment analysis identified nutrient reservoir activity as the most significant term among differentially expressed genes (Supplementary Figure S2C).

To test whether *rgh6* affects alternative splicing, we used rMATS (Shen et al. 2012; Shen et al. 2014) to compare normalized read counts mapped to exon-exon junctions for inclusion or exclusion isoforms to estimate the exon inclusion level. For a given exon, the inclusion level was calculated by the percent of exon inclusion reads of the total junction reads for both inclusion and exclusion events (Figure 6D). Although rMATS identifies all major types of alternative splicing, including exon skipping (SE), alternative 5’ splice sites (A5SS), alternative 3’ splice sites (A3SS), mutually exclusive exons (MXE), and retained introns (RI), we focused on RI events because these are the most common form of alternative splicing regulation in plants (Ner-Gaon et al. 2004; Filichkin et al. 2010; Marquez et al. 2012).

A total of 11,959 RI events were scored, in which 301 events were statistically significant with a false discovery rate (FDR) <0.05. We overlapped these RI events with differentially expressed genes (DEGs), of which 154 events remained. Only 24 RI events had a mean FPKM of ≥10 in both *rgh6* mutant and normal siblings to ensure sufficient read depth for accurate splice junction detection (Figure 6E). We surveyed these 24 RI events by the inclusion level, but none of them had large magnitude effects on intron splicing. We further selected two out of the 24 RI events and visualized them with the Integrative Genomics Viewer (IGV) (Robinson et al. 2011). Although the two events are differentially expressed, differential splicing events represented a minor fraction of the total expression levels suggesting RI splicing had little impact on differential expression or the expressed isoforms of these two genes between normal and *rgh6* (Figure 6F and 6G).

Both *rgh3* and *rbm48* are maize *rgh* mutants that encode minor intron RNA splicing factors that cause U12-type mis-splicing events (Gault et al. 2017; Bai et al. 2019). To further test whether *rgh6* affects minor intron splicing, six genes with U12-type introns and shown to be mis-spliced in *rgh3* and *rbm48* were selected for RT-PCR assays that included the U12-type introns. These assays showed no evidence for minor intron splicing defects in the *rgh6* mutant (Supplementary Figure S3). Combined, these results suggest that *rgh6* is unlikely to affect mRNA splicing at a genome-wide level.

### Aberrant miRNA processing in *rgh6* mutants

Six miRNAs are highly expressed across maize tissues at >4,000 average mapped reads per million (RPM) and include zma-MIR156b, zma-MIR159a, zma-MIR167h, zma-MIR168b, zma-MIR171j, zma-MIR319b (Zhang et al. 2009). For 10 DAP endosperm, all six mature miRNAs were at lower levels in *rgh6* mutants than in normal siblings (Figure 7A). These data suggest that *rgh6* has global defects in miRNA accumulation. Primers amplifying both pri-miRNA and pre-miRNA for these six miRNAs showed higher levels of these precursor species in *rgh6* compared with normal siblings (Figure 7B). An accumulation of precursor and reduction of product RNA species is consistent with *rgh6* functioning in miRNA processing. It is possible that *rgh6* has an indirect effect on miRNA accumulation by influencing expression of the miRNA dicing complex subunits, DCL1, HYL1, and SERT1-SERT4. However, transcript levels for all six dicing complex subunits are increased in *rgh6* mutant relative to normal siblings (Figure 7C).

**Figure 7.**
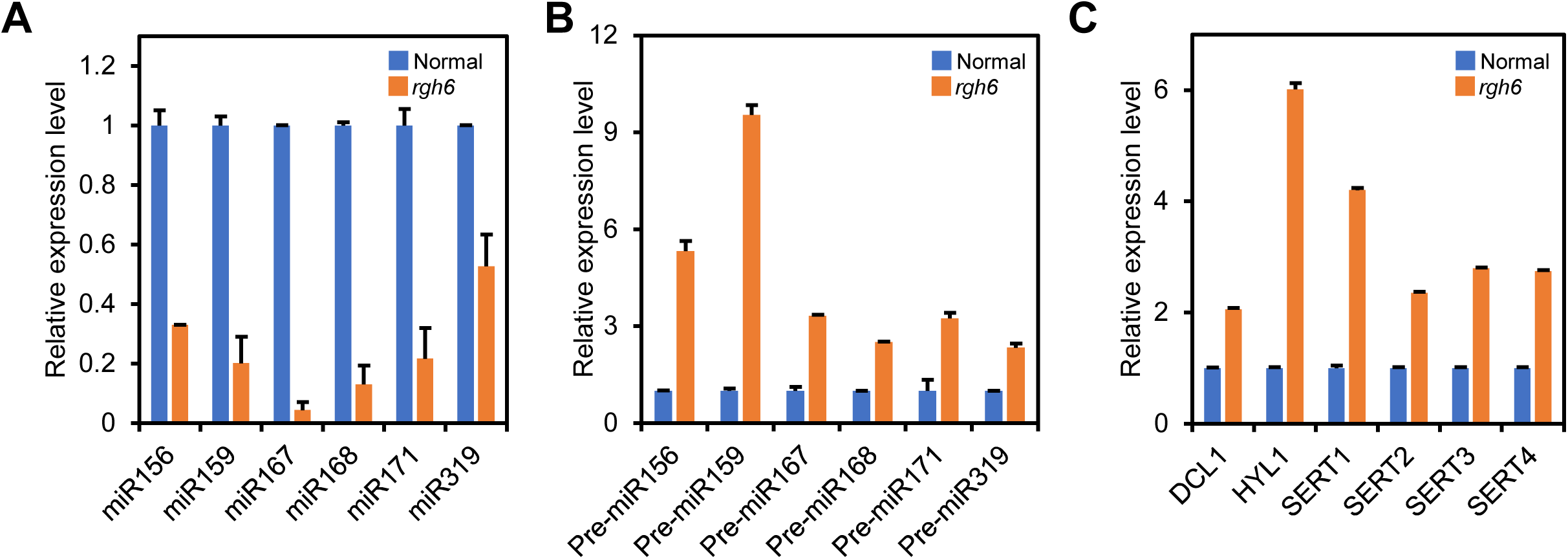
*rgh6* is required for miRNA processing. (A-C) RT-qPCR analysis of (A) mature miRNA, (B) pri-/pre-miRNA, and (C) Dicing complex proteins in 10 DAP endosperm. Graphs plot the average of three biological replicates from independent developing ears and four technical replicates of each biological replicate. Error bars are standard deviation of the three biological replicates. Expression is relative to normal samples for each PCR product. Control genes were U6 mature miRNA and *Actin1* for pri-/pre-miRNA and mRNA.

In plants, miRNA predominantly impact gene expression through targeted degradation of mRNA. We surveyed the global expression profile of predicted miRNA targets in 10 DAP endosperm mRNA-seq data. A total of 247 miRNA targeted genes were predicted based on small RNA-seq from five B73 maize tissues (Zhang et al. 2009). Of these predicted targets, 31 were found to be differentially expressed in *rgh6* with 20 genes having higher expression and 11 genes having lower expression in mutant endosperm tissue. Compared to the entire set of *rgh6* DEGs with 2,270 increased and 1,176 decreased transcripts in *rgh6*, the differentially regulated miRNA targets had a similar ratio of up and down regulated genes suggesting that reduction of multiple miRNA levels does not result in a general increase of miRNA target transcripts (Supplementary Table S1).

### Mutation in *rgh6* disrupts endosperm cell differentiation

In the endosperm, *Rgh6* is predominantly expressed from 6 to 8 DAP, which is coincident with the proliferation-differentiation stage of development (Chen et al. 2014; Yi et al. 2019) (Supplementary Figure S4A and S4B). An RNA-seq analysis of laser-capture micro-dissected maize kernels at 8 DAP showed that *Rgh6* transcripts can be detected in endosperm cell types with the highest signal in aleurone layer (Zhan et al. 2015) (Supplementary Figure S4C). Transcriptome data from hand dissected maize kernels at 13 DAP showed that *Rgh6* transcripts can be detected in embryo/endosperm interfaces with the highest signal in the EAS endosperm tissue (Doll et al. 2020) (Supplementary Figure S4D). These published data show that *Rgh6* transcripts are highly expressed in endosperm epidermal cell types.

To test the developmental consequences of *rgh6* mutation, we surveyed the FPKM values of genes encoding starch biosynthesis enzymes and seed storage proteins from the *rgh6* endosperm mRNA-seq data. Among 16 genes known to be part of the starch and zein biosynthetic pathways, transcript levels of all were reduced by at least 50% in the *rgh6* mutant. The 19 kDa α-zein and 22 kDa α-zein were reduced more than 90% (Supplementary Figure S5; Supplementary Figure S6). These data suggest that *rgh6* mutants reduce starch and storage protein accumulation in the endosperm.

We also analyzed transcript levels of several endosperm cell-type marker genes in *rgh6* mutant kernels at 10, 12, and 14 DAP, including the *az22z5* α-zein gene specific to the starchy endosperm (Woo et al. 2001), *Al9* specific to the aleurone layer (Gómez et al. 2009), *Esr1* for the ESR (Opsahl-Ferstad et al. 1997), the *Betl2* and *Tcrr1* markers for the BETL, (Hueros et al. 1999; Muñiz et al. 2006) and *Sweet15a* specific to the EAS (Doll et al. 2020). Compared with normal kernels, *rgh6* kernels showed decreased expression of the starchy endosperm cell marker, *az22z5*, while expression of the endosperm epidermal cell markers, *Al9*, *Esr1*, *Betl2*, *Tcrr1* and *Sweet15a,* was increased (Figure 8). These results indicated that *rgh6* disrupts endosperm cell type patterning.

**Figure 8.**
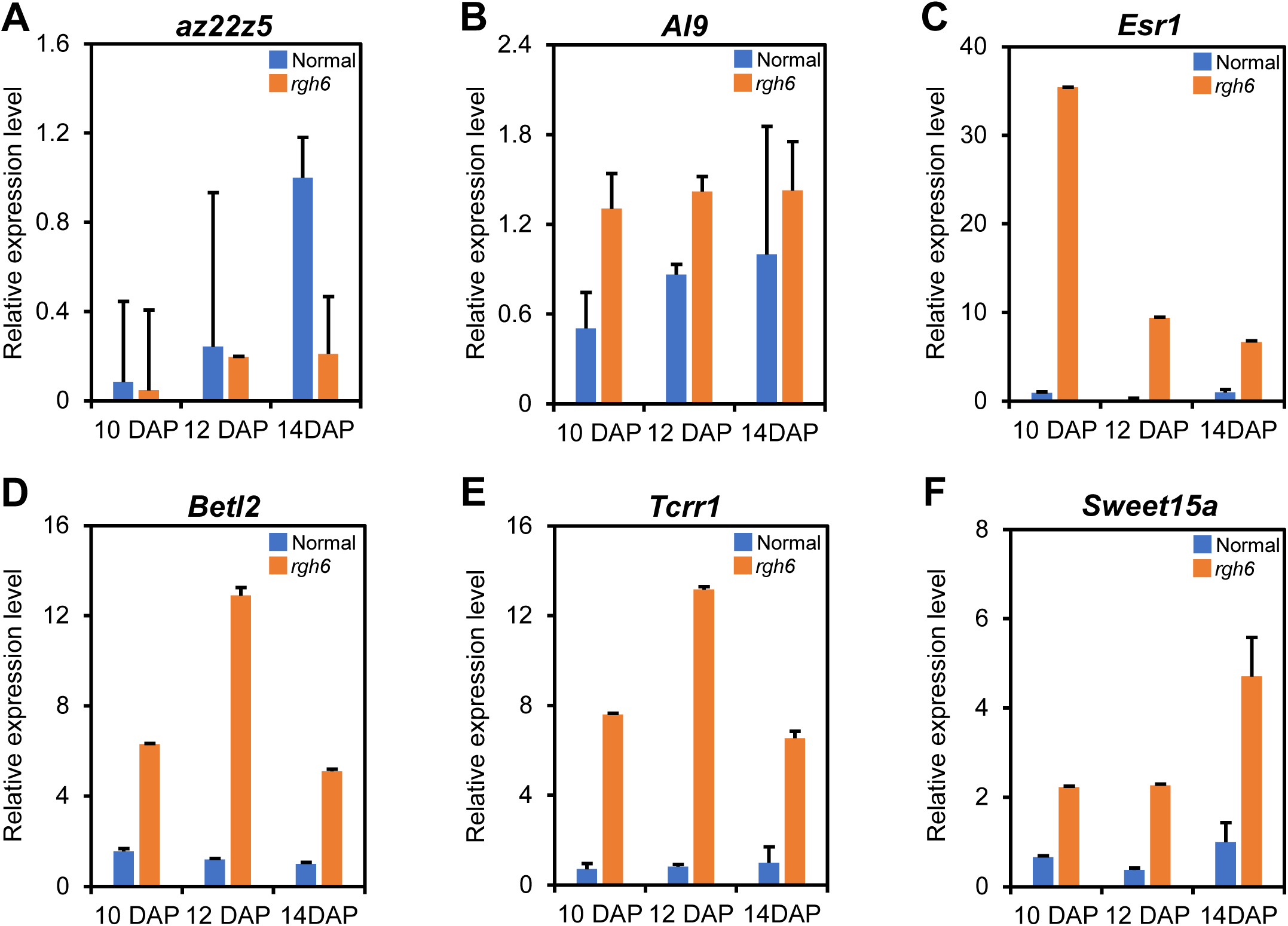
*rgh6* kernels have increased levels of endosperm epidermal cell markers. (A-F) RT-qPCR analysis of endosperm cell type marker genes in total RNA extracted from whole kernels at 10, 12, and 14 DAP. Graphs plot averages of three biological replicates from independent ears and four technical replicates. Error bars are standard deviation of three biological replicates. Expression is relative to 14 DAP normal samples in all panels. *Actin1* was used as the internal control.

### *rgh6* impacts on known developmental miRNA interactions

To further investigate the impact of aberrant miRNA processing on endosperm cell patterning in *rgh6* mutant, we focused on DEGs that are predicted miRNA target genes and have reported biological functions in maize (Supplementary Table S2; Supplementary Table S3). The predicted target of miR528, *bige1*, impacts lateral organ number and was also downregulated in *rgh6* (Suzuki et al. 2015). Although *bige1* is expressed during normal endosperm development, no endosperm phenotypes have been reported for mutant alleles. In addition, *myb138* is a verified target of miR159 and over expression of miR159 increases maize endosperm cell division and grain size (Wang et al. 2023). We observed reduced *myb138* transcript levels in *rgh6* consistent with the smaller grain of mutant kernels.

Developmental phase change is regulated by expression levels of mature miR156 with higher levels of miR156 prolonging the juvenile state (Zhao et al. 2023). In *rgh6* endosperm, there are two predicted targets of miR156 that are up-regulated DEGs. One, *sbp22*, is an SPL transcription factor that has a function in flowering time (Jin et al. 2023). The second miR156 target is a predicted histidine phosphotransfer protein that is predominantly expressed in aleurone and proliferating cells at 6-7 DAP (Yuan et al. 2024). Similarly, the NAC transcription factor, Zm00001d016950, is up-regulated in *rgh6*. This gene is predominantly expressed in aleurone cells at 6-7 DAP and is a predicted target of miR164. Increase of miR156 targets and precocious expression of differentiated epidermal endosperm cell type markers in *rgh6* mutants would be consistent with epidermal endosperm cells advancing through developmental phases more rapidly.

Increased levels of miR172 microRNA induces flowering in sporophytes but does not have a known role in endosperm development (Zhao et al. 2023). One miR172 target, *ts6*, functions in inflorescence development and had increased transcript levels in *rgh6* (Chuck et al. 1998; Chuck et al. 2008). Another miR172 predicted target, Zm00001d026448, encodes an AP2/EREBP transcription factor and had lower transcript levels in *rgh6*.

Although the remaining miRNA predicted targets do not have experimentally validated roles in maize, multiple transcription factors that are predicted targets of miR160, miR166, miR167, miR396, and miR482 suggest *rgh6* endosperm may have more of a stem cell-like fate. Predicted targets of miR160 and miR167 include five Auxin Response Factor genes that are upregulated suggesting increased sensitivity to auxin in *rgh6*. For miR396, three Growth Regulating Factor genes are increased in *rgh6*, which would be expected to promote cell proliferation. Zm00001d025694 is an upregulated SURF2 homolog that is a predicted target of miR482. In humans, SURF2 is associated with cancer and proliferating cells by promoting ribosome synthesis (Tagnères et al. 2024), while the maize gene is expressed predominantly in proliferating endosperm cells at 6-7 DAP (Yuan et al. 2024). Finally, miR166 restricts expression of *HD-ZIPIII* genes, which specify adaxial, endodermal, and meristematic fates (Juarez et al. 2004; Kidner and Martienssen, 2004; Nogueira et al. 2007; Chitwood et al. 2009; Carlsbecker et al. 2010; Miyashima et al. 2011). In *rgh6* endosperm, there are two up-regulated predicted targets of miR166, including an HD-ZIP III gene (Zm00001d031061), which may be consistent with a greater stem cell-like fate in the mutant.

## Discussion

### *rgh6/dek51* is unlikely to affect rRNA and mRNA processing

We report an independent identification of the *rgh6/dek51* locus and characterization of *rgh6* function in the endosperm. DEAD-box RNA helicases have diverse functions in plant RNA processing (Li et al. 2023). It is critical to consider multiple lines of evidence before proposing a specific RNA processing function for RGH6. Consistent with protein sequence analysis predictions (Kosugi et al. 2008; Kosugi et al. 2009 a; Kosugi et al. 2009 b), we found that a RGH6-GFP fusion protein localizes within the nucleus in the nucleolus and punctate loci that are potentially nuclear speckles. This subcellular localization indicates a function in processing nuclear RNA transcripts such as pre-rRNA, pre-mRNA, pri-miRNA, or other non-coding RNA species.

Wang et al. (2024) suggested that RGH6/DEK51 has a primary function in 3’ end processing of pre-rRNA despite their observation that mature 25S, 18S, and 5.8S levels are not changed between mutant and normal embryos. In agreement, we also found that the levels of 25S and 18S are statistically equivalent in *rgh6* mutants and normal sibling endosperm tissues. Wang et al. (2024) showed subtle differences in levels of 45S precursor and pre-rRNA intermediates in *dek51* mutant embryos. By contrast, mutations in Arabidopsis DEAD-box RNA helicases that are required for rRNA processing show lethal transmission defects through the female gametophyte. The Arabidopsis SLOWWALKER3 (SWA3) locus encodes the RH36 DEAD-box protein (Huang et al. 2010; Liu et al. 2010). Mutation of *RH36* results in reduced female transmission, delayed female gametophytic cell cycle, and increased unprocessed 18S pre-rRNA accumulation. Another Arabidopsis DEAD-box RNA helicase, RH29, is required for both male and female gametophyte development and can be partially rescued by its yeast homolog Dbp10p, which has a demonstrated rRNA processing role in yeast (Chen et al. 2020). We did not observe gametophyte or transmission defects in *rgh6* mutants. Based on these data, *rgh6* is unlikely to impact rRNA processing sufficiently to cause the defective endosperm and embryo phenotypes observed.

DEAD-box RNA helicases also function in mRNA processing. The homologous Arabidopsis RCF1 and rice RH42 DEAD-box proteins are critical for pre-mRNA splicing under cold stress (Guan et al. 2013; Lu et al. 2020). RCF1 is localized in the nucleus, and many cold-responsive genes are mis-spliced in the *rcf1-1* mutant under cold treatment (Guan et al. 2013). Both knock-down and over-expression of rice *RH42* confer hypersensitivity to cold stress, and a genome-wide survey of alternative splicing under cold treatment showed that RH42 promotes intron removal for a subset of cold-induced transcripts (Lu et al. 2020).

Our *rgh6* transcriptome analysis did not reveal mRNA splicing defects, and we are not aware of any defective seed mutants encoding DEAD-box helicases and required for mRNA splicing. However, two maize *rgh* mutants, *rgh3* and *rbm48*, encode minor intron RNA splicing factors and show similar subnuclear localization patterns to RGH6 (Fouquet et al. 2011; Gault et al. 2017; Bai et al. 2019). A directed test of genes with minor, U12-type introns found no minor intron splicing defects in *rgh6* mutants. From these data, we conclude that *rgh6* is unlikely to affect mRNA processing.

### *rgh6* is required for miRNA processing

Multiple RNA helicases are known to affect miRNA processing, such as CHR2 (Wang et al. 2018), SMA1 (Li et al. 2018), and RH6/RH8/RH12 (Li et al. 2021). Plant miRNAs biogenesis has three major steps within the nucleus: transcription, dicing complex cleavage, and 3’ methylation. Mutations that impact transcript production, such as *sma1* in Arabidopsis (Li et al. 2018), have reduced levels of both mature and pri-miRNA. Arabidopsis mutants in dicing complex subunits accumulate increased levels of pri-miRNA concomitant with reduced miRNA processing (Yang et al. 2006). Similarly, mutants that impact the processing of pri-miRNA outside of the D-body, such as *mos2* in Arabidopsis (Wu et al. 2013), have decreased levels of mature miRNA but increased levels of pri-miRNA.

Among Arabidopsis genes, *Rgh6* is most closely related to *RH27* (Hou et al. 2021). Both *rgh6* and *rh27* mutants share similarities with both loci causing zygotic lethal seed defects and a decrease in mature miRNA. However, the levels of pri-miRNA and mature miRNA in *rh27* mutants are reduced suggesting either a direct role in pri-miRNA transcription or a co-transcriptional process required for pri-mRNA transcript accumulation. By contrast, *rgh6* has increased levels of pri-miRNA/pre-miRNA and decreased levels of miRNA, suggesting that RGH6 acts after *MIR* gene transcription. It is also possible that the differences in pri-miRNA/pre-miRNA levels observed in *rgh6* and *rh27* mutants are due to tissue specific differences in miRNA regulatory pathways in *rgh6* endosperm versus *rh27*shoot apices and root tips.

Our genome-wide analysis of predicted miRNA targets in the *rgh6* endosperm showed that 65% of differentially expressed target genes have higher transcript levels relative to normal siblings, which is consistent with genome-wide DEG trends for the mutant. Although miRNA-directed cleavage of target mRNAs is the predominant mechanism of target inhibition in plants, studies of several Arabidopsis miRNA processing mutants also found reduced transcript levels of a subset of target mRNAs (Li et al. 2020; Hou et al. 2021; Xu et al. 2023). Intriguingly, miRNA target genes show contrasting trends in *rh27* transcriptome data with 63% of predicted targets increased in shoot apices and only 33% of predicted targets increased in root tips (Hou et al. 2021). Thus, reduced miRNA levels have tissue-specific consequences on target gene transcript levels.

### Dysregulation of miRNA processing disrupts endosperm cell differentiation

In maize, modulation of miR169o and miR159 levels affects seed size by regulating transcription factors involved in endosperm cell division (Zhang et al. 2022; Wang et al. 2023). Transcript levels for one of these transcription factor targets, *Myb138*, are reduced in *rgh6* and is consistent with the reduced grain-fill in mutants. However, *rgh6* shows more complex dysregulation of miRNA target genes that suggests miRNA participate in endosperm cell fate decisions and the timing of endosperm development. Low levels of miR156 and the significant increase in transcript levels for predicted miR156 targets suggest some aspects of developmental phase change are impacted. The precocious expression of endosperm markers for aleurone, ESR, BETL, and EAS, which are all epidermal cell fates, contrasts with down regulation of starchy endosperm specific genes *rgh6*. These cell markers suggest reduced miRNA promotes epidermal cell differentiation. However, multiple miRNA target genes, such as ARF and GRF transcription factors, are up regulated that suggest an increase in stem or proliferating cells in *rgh6*. Potentially, global reduction in miRNA prolongs a stem cell-like fate for internal endosperm, while promoting rapid differentiation of epidermal fates.

In conclusion, our study identified a DEAD-box RNA helicase required for normal miRNA processing. Dysregulation of miRNA target genes and endosperm cell type marker genes in *rgh6* support the hypothesis that miRNA regulation is required for normal endosperm cell patterning. Genetic chimeras further indicate that the loss of normal endosperm cell patterning negatively affects embryo development leading to early embryo developmental arrest.

## Materials and methods

### Genetic stocks

Maize was grown in the field at the University of Florida Plant Science Research and Education Unit in Citra, FL and in the greenhouse at the Horticultural Sciences Department in Gainesville, FL. The *rgh6-umu1 and rgh*-744* alleles were isolated from the UniformMu forward genetics screen (McCarty et al. 2005). The *rgh6-umu2* allele was identified from the UniformMu reverse genetics resources as insertion mu1095038::Mu in stock UFMu -11204 (Settles et al. 2004; Settles et al. 2007). All mutants were maintained in the color converted W22 inbred background.

### Morphology

Ears and kernels were imaged on a flatbed scanner. Ears segregating for *rgh6* in the W22 background and the B73/W22 hybrid background were sorted, counted, and calculated between normal and *rgh* kernels. For mature kernel microscopy, 15 normal and 15 *rgh* kernels were sampled from three segregating ears and soaked overnight in sterilized water. Half-kernel sagittal sections, cut with a utility knife, were split for imaging and genotyping by PCR.

For germination experiments, 15 normal and 15 *rgh* kernels from three segregating ears were rolled tightly in paper towels rinsed with sterile water and placed in a 2 L glass beaker with 0.5 L sterilized water. The beaker with paper rolls were moved to the greenhouse. Germination frequency and seeding phenotypes were recorded 7 days after sowing.

### B-A translocation

The *rgh*-744* was mapped by bulk segregant analysis and used as a control for *rgh6* B-A translocation genetics. Pollen from TB-5Sc hyperploid plants was crossed onto 10 to 15 plants segregating for *rgh6/+* or *rgh*-744/+* (Birchler and Alfenito, 1993). Non-concordant kernels were selected from uncovered F_1_ ears, sectioned with a utility knife, and imaged on a flatbed scanner.

### Molecular cloning of *rgh6*

The *rgh6-umu1* allele was mapped from a *rgh6*/+ x B73 F_2_ mapping population. DNA was extracted from individual F_2_ mutant kernels and genotyped with SSR or insertion-deletion polymorphism (IDP) markers (Martin et al. 2010; Settles et al. 2014). Two polymorphic markers per chromosome arm were used to scan the genome using 24 *rgh* kernels and a normal sibling pool. After detecting linkage for umc1587, UFIDP5-3.72 and IDP5934 were identified as flanking markers for the initial mapping population. Fine mapping used 244 *rgh* kernels that were genotyped with four IDP markers, UFIDP5-3.72, UFIDP5-3.77, UFIDP5-3.83, and UFIDP5-4.15. *Mu* flanking sequences from the *rgh6* alleles were amplified, purified, and blasted against GenBank maize sequences (Settles et al. 2004). Co-segregation of the *rgh6-umu1* allele with the mutant phenotype was confirmed by amplifying normal or *Mu*-insertion sequences using Rgh6-F1, Rgh6-R1, and TIR6 primers from a segregating self-pollination in the UniformMu genetic background (Settles et al. 2007). The *rgh6-umu2* allele (mu1095038::Mu) was confirmed by reciprocal crosses between the UFMu-11204 stock and *rgh6-umu1* stocks. First ears were crossed to test genetic complementation, and second ears were self-pollinated to confirm plant genotypes. Primer sequences for molecular mapping are listed (Supplementary Table S4).

### Subcellular localization

RNA was extracted from W22 kernels using TRIzol and DNase I (Invitrogen). The full-length CDS were amplified from W22 kernel cDNA with Phusion high-fidelity DNA polymerase (New England Biolabs). The PCR products were subcloned into the pENTR/D-TOPO vector (Invitrogen) and then recombined in the binary vector pGWB504 (Karimi et al. 2002; Karimi et al. 2007). The binary constructs were transformed into the *Agrobacterium tumefaciens* strain GV3101 with a freeze-thaw method (Wise et al. 2006). Agrobacteria carrying individual constructs were mixed in a 1:1 ratio before infiltration. The abaxial surface of leaves from 4- to 6-week-old *N. benthamiana* plants were infiltrated with a needleless syringe. The fusion proteins were visualized 48 to 72 h after transient transformation (Kapila et al. 1997). A 100 ng/mL DAPI solution, which used for staining the nucleus, was injected to the leaf abaxial surface. Approximately 1 cm^2^ areas of leaf abaxial surface were embedded on the slides and rinsed with distilled water. The slides were incubated in the dark for 3 to 5 minutes prior to imaging. The fluorescent signals were captured under a Zeiss LSM 710 confocal microscope. The GFP was excited at 488 nm and detected with an emission band of 522-572 nm. The mCherry was excited at 587 nm and detected with an emission band of 590-630 nm. The total number of leaves infiltrated and imaged for each experimental condition were five leaves for Free-GFP, five leaves for RGH6-GFP, and eight leaves for RGH6-GFP and PRH75-mCherry co-localization. Transient expression experiments were conducted in three replicate procedures with similar results. Primer sequences for transient expression are listed (Supplementary Table S4).

### RNA extraction

Developing kernels of self-pollinated *rgh6-umu2* heterozygotes were sampled at 10, 12, and 14 days after pollination (DAP). RNA was extracted from both whole kernels and dissected endosperm from three independent ears. Tissues were frozen in liquid nitrogen and ground into fine powder with a mortar and pestle. 100 mg ground tissue was mixed with 200 µL RNA extraction buffer (50 mM Tris-HCl, pH 8, 150 mM LiCl, 5 mM EDTA, 1% SDS in DEPC-treated water). RNA was first extracted with phenol: chloroform and TRIzol, then precipitated with isopropanol and washed with 70% ethanol, and finally resuspended in nuclease-free water treated with DNase I. RNA was further purified using a RNeasy MinElute Cleanup Kit (Qiagen) and stored at -80 °C.

### RT-PCR and RT-qPCR

Total RNA (1 µg) extracted from 10 DAP endosperm was reverse transcribed to cDNA with M-MLV reverse transcriptase (Promega, Catalog # M170A) and Oligo (dT)_15_ (Promega, Catalog # C110A). Specific RT-PCR reaction conditions were optimized for cycle number and annealing temperature for each primer pair.

Self-pollinated ears segregating for *rgh6-umu2* allele were sampled at 10, 12, and 14 DAP. RNA isolated from whole kernels at three time points was used for cell marker analysis. RNA isolated from endosperm tissues at 10 DAP was used for mature miRNA and precursor miRNA analysis. The expression level of cell markers was analyzed with the Power SYBR Green PCR Master Mix (Applied Biosystems). Primer sequences for endosperm cell marker are listed (Supplementary Table S4). The expression level of mature miRNA was analyzed with the Mir-X miRNA qRT-PCR SYBR Kit (Takara). The expression level of miRNA precursor was analyzed with the Power SYBR Green PCR Master Mix (Applied Biosystems). Primer sequences for miRNA processing analysis are listed (Supplementary Table S4). All RT-qPCR reactions were completed with a Bio-Rad CFX96 Touch Real-Time PCR Detection System. Normalization was relative to *Actin 1* for cell marker and precursor miRNA, and U6 snRNA for miRNA. The relative expression was calculated by the comparative cycle threshold (2^-ΔΔCt^) method (Livak and Schmittgen, 2001). The relative expression was the average of three technical replicates of three biological endosperm pools from independent segregating ears.

### rRNA and mRNA analyses

The quality and quantity of endosperm RNA samples were confirmed by Bioanalyzer and Qubit. RNA isolated from endosperm tissues for paired mutant and normal samples was used. RNA was initially visualized by agarose gel electrophoreses, and 200 ng of RNA per sample was assayed by Agilent 2100 Bioanalyzer.

Six poly(A) captured, non-directional cDNA libraries were constructed for 10 DAP normal and *rgh6-umu2* endosperm and sequenced on a Illumina Novaseq 6000 platform with 150 bp paired-end reads. Raw reads were processed by removing 5’ and 3’ adaptors, poly(N) segments and low-quality reads. Clean reads were aligned to B73 RefGen_v4 maize genome assembly with Hisat2 v2.0.5 (Kim et al. 2015; Pertea et al. 2016). Read counts were calculated by featureCounts v1.5.0-p3 (Liao et al. 2014), and transcript levels was normalized to FPKM. DEGs were identified with the DESeq2 R package (1.20.0) with the criteria of |log2 fold change|≥1 and adjusted P-values <0.05 (Love et al. 2014). The R package, clusterProfiler, was used to identify enriched Gene Ontology terms within the DEGs (Yu et al. 2012).

Global mRNA splicing was analyzed with rMATS using normalized read counts mapped to every exon-exon junction for calculation of exon inclusion levels (Shen et al. 2012; Shen et al. 2014). Briefly, rMATS models a multivariate uniform prior for the overall similarity of alternative splicing profiles between samples based on exon inclusion levels for all alternatively spliced exons. A Bayesian posterior probability for differential alternative splicing was calculated using a Markov chain Monte Carlo (MAMC), and rMATS calculated a p-value for each exon. The p-values were corrected for FDR using the Benjamini and Hochberg’s method. RI events identified were visualized with the Integrative Genomics Viewer (Robinson et al. 2011).

### miRNA processing analysis

Maize miRNA genes were grouped into families based on mature miRNA sequences (Zhang et al. 2009). Expression levels of each family was averaged based on the RPM of root, seeding, tassel, ear, and pollen to identify miRNA families with broad expression across tissues and tested for endosperm expression of pri/pri-miRNA and miRNA species by RT-qPCR.

Maize miRNA target gene identifiers were converted from B73 RefGen_v1 to B73 RefGen_v4 maize genome assembly (Zhang et al. 2009). The rgh6 DEGs were identified with the DESeq2 R package (1.20.0) with the criteria of fold change ≥2 and adjusted p-value <0.05 (Love et al. 2014) (Supplementary Table S2). Differentially expressed targets were the overlap between *rgh6* DEGs and maize miRNA targets (Supplementary Table S3).

## Acknowledgements

We thank Fang Bai, Chiwah Tseung, Noriko Suzuki, and David Silverman at the University of Florida for technical assistance.

## Author contributions

L.C.H. and A.M.S. designed the research. T.Y. and M.S. performed the experiments. T.Y. analyzed the data. T.Y. drafted the manuscript. M.S., L.C.H. and A.M.S. revised the manuscript. All authors approved the final version of the manuscript.

## Supplementary data

The following materials are available in the online version of this article.

Supplementary Figure S1. Mapping strategy of *rgh6* mutant.

Supplementary Figure S2. mRNA sequencing of *rgh6* mutant.

Supplementary Figure S3. Analysis of U12-type intron splicing.

Supplementary Figure S4. *Rgh6* transcript levels from published transcriptome studies.

Supplementary Figure S5. Transcript levels of starch synthesis genes in *rgh6* endosperm.

Supplementary Figure S6. Transcript levels of zein seed storage protein related genes in *rgh6* endosperm.

Supplementary Table S1. Comparison of transcriptome and miRNA target gene expression differences in mutants affecting miRNA processing.

Supplementary Table S2. Differentially expressed genes identified in maize *rgh6* mutants relative to normal siblings.

Supplementary Table S3. Differentially expressed miRNA target genes identified in maize *rgh6* mutants relative to normal siblings.

Supplementary Table S4. Primers used in this study.

## Funding

This work was supported by the Vasil-Monsanto Endowment of Plant Cell and Molecular Biology to A.M.S and the Graduate School Preeminence Award of University of Florida to T.Y.

## Data availability

Raw data from RNA-seq has been deposited in the NCBI-SRA (https://www.ncbi.nlm.nih.gov/sra) under the accession number PRJNA1190850.

## Conflict of interest

The authors declare no competing interests.

## Supplementary Figure legends

**Supplementary Figure S1. Mapping strategy of *rgh6* mutant. (A)** Schematic of the F_2_ mapping strategy. Red rectangle indicates *rgh* kernels used for molecular mapping in the F2 population. **(B)** Self-pollinated ear segregating for *rgh6* in the B73/W22 hybrid background. Arrowheads indicate *rgh* kernels. Scale bar = 1 cm. **(C)** Segregation of *rgh* phenotypes in four independent self-pollinated *rgh6* heterozygotes in the B73/W22 hybrid background.

**Supplementary Figure S2. mRNA sequencing of *rgh6* mutant. (A)** PCA plot of differentially expressed genes (DEGs) between *rgh6* and its normal siblings, respectively. Red and blue dots are normal and mutant samples. Endosperm tissues at 10 DAP were dissected from normal and *rgh6* kernels. Total RNAs were extracted from three biological replicates of paired *rgh6* mutant and normal sibling pools. Each replicate is a self-pollinated ear segregating for *rgh6* in W22 background. **(B)** Volcano plot of DEGs between *rgh6* and normal siblings. Red dots are up-regulated genes in the mutant versus normal. Green dots are the down-regulated genes in the mutant versus normal. Blue dots are genes without a significant difference. **(C)** Bar plot for GO enrichment of DEGs. The red bars indicate GO terms for biological process. The green bars indicate GO terms for cellular component. The blue bars indicate GO terms for molecular function. Values on bars are the number of DEGs for each GO term.

**Supplementary Figure S3. Analysis of U12-type intron splicing. (A)** Schematics show amplified products with PCR primers indicated by arrows. **(B)** RT-PCR of *rgh6* mutant (M) and normal siblings (N) RNA from endosperm tissue with three biological replicates at three developmental time points.

**Supplementary Figure S4. *Rgh6* transcript levels from published transcriptome studies. (A-B)** Temporal expression of *Rgh6* locus in embryo **(A)** and endosperm **(B)**. FPKM values are from Chen et al. (2014). **(C)** Spatial expression of *Rgh6* locus from dissected seed tissues at 8 DAP. FPKM values are from Zhan et al. (2015). **(D)** Spatial expression of *Rgh6* locus from dissected seed tissues at 13 DAP. FPKM values are from Doll et al. (2020).

**Supplementary Figure S5. Transcript levels of starch synthesis genes in *rgh6* endosperm. (A-H)** Mean and standard deviation FPKM values are plotted from the three biological replicates of 10 DAP endosperm RNA-seq dataset for *rgh6* and normal siblings.

**Supplementary Figure S6. Transcript levels of zein seed storage protein related genes in *rgh6* endosperm. (A-H)** Mean and standard deviation FPKM values are plotted from the three biological replicates of 10 DAP endosperm RNA-seq dataset for *rgh6* and normal siblings.

## Notes

### Competing Interest Statement

The authors have declared no competing interest.

