## Supplemental Figures for "Maize *rough endosperm6* is a predicted RNA helicase required for miRNA processing and endosperm cell patterning"

A

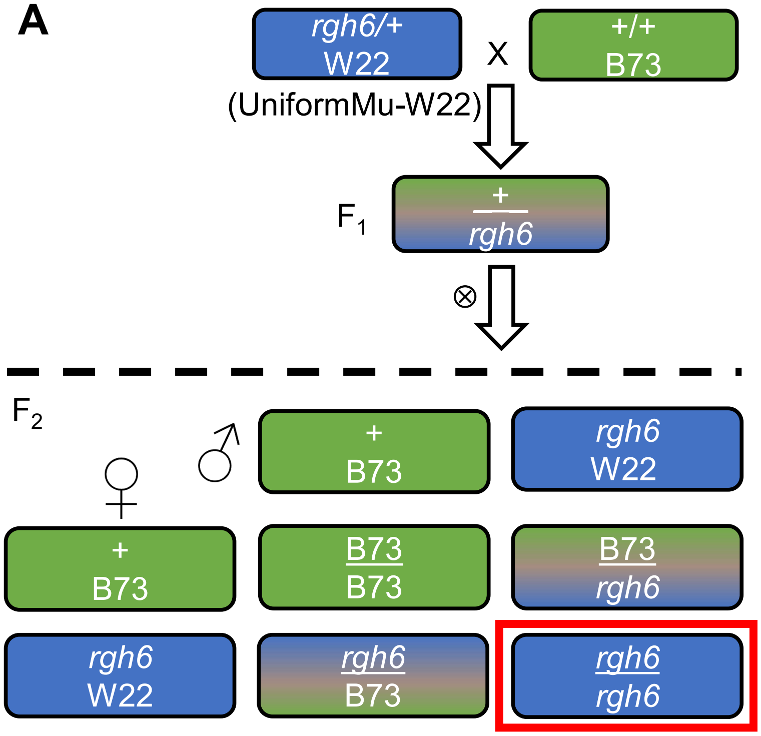


B

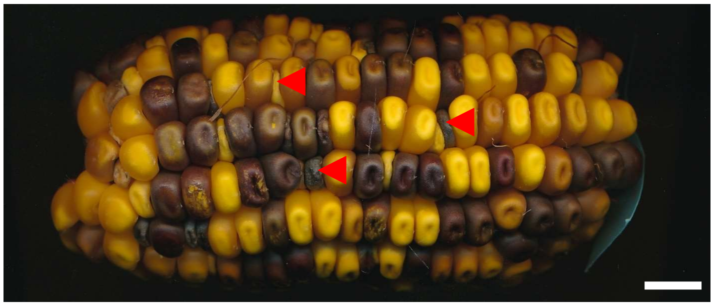

*rgh6*/+ ⊗

C

| Pollination | Normal | *rgh* | Ratio | Expected ratio | p-value ($X^2$ for 3:1) |
|---|---|---|---|---|---|
| Ear1_*rgh6*/+⊗ | 326 | 101 | 3.23:1 | 3:1 | 0.520 |
| Ear2_*rgh6*/+⊗ | 399 | 123 | 3.24:1 | 3:1 | 0.448 |
| Ear3_*rgh6*/+⊗ | 322 | 107 | 3.01:1 | 3:1 | 0.978 |
| Ear4_*rgh6*/+⊗ | 213 | 62 | 3.44:1 | 3:1 | 0.347 |

**Supplementary Figure S1. Mapping strategy of *rgh6* mutant. (A)** Schematic of the $F_2$ mapping strategy. Red rectangle indicates *rgh* kernels used for molecular mapping in the F2 population. **(B)** Self-pollinated ear segregating for *rgh6* in the B73/W22 hybrid background. Arrowheads indicate *rgh* kernels. Scale bar = 1 cm. **(C)** Segregation of *rgh* phenotypes in four independent self-pollinated *rgh6* heterozygotes in the B73/W22 hybrid background.

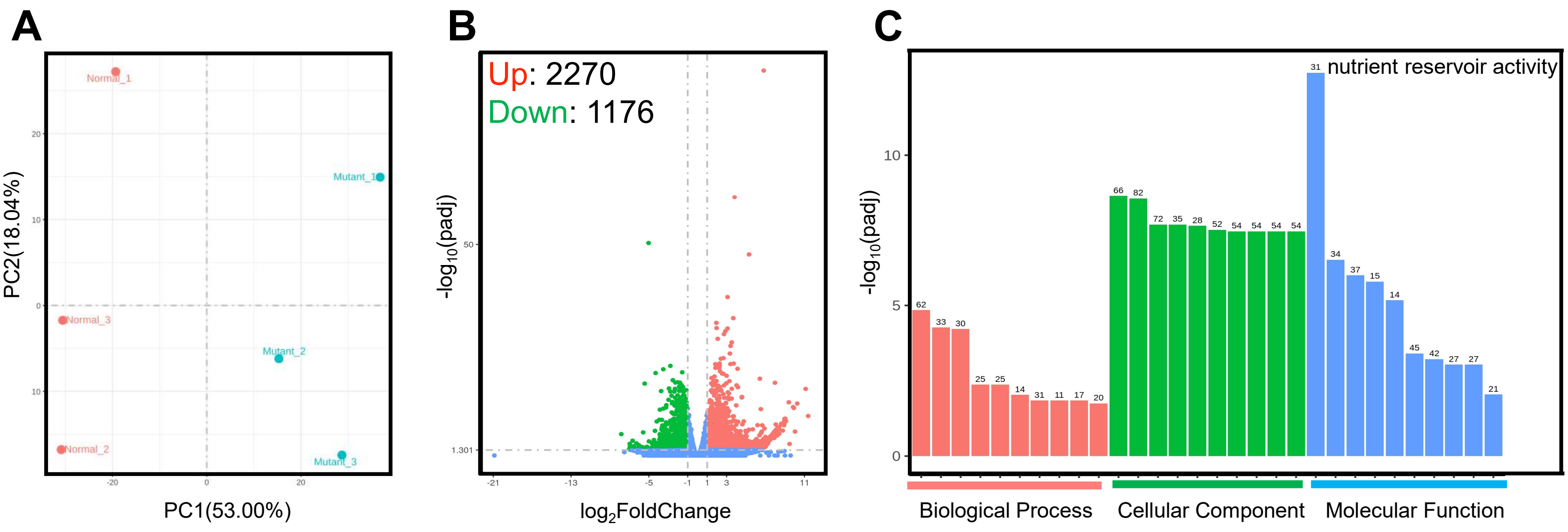


**Supplementary Figure S2. mRNA sequencing of *rgh6* mutant. (A)** PCA plot of differentially expressed genes (DEGs) between *rgh6* and its normal siblings, respectively. Red and blue dots are normal and mutant samples. Endosperm tissues at 10 DAP were dissected from normal and *rgh6* kernels. Total RNAs were extracted from three biological replicates of paired *rgh6* mutant and normal sibling pools.  Each replicate is a self-pollinated ear segregating for *rgh6* in W22 background. **(B)** Volcano plot of DEGs between *rgh6* and normal siblings. Red dots are up-regulated genes in the mutant versus normal. Green dots are the down-regulated genes in the mutant versus normal. Blue dots are genes without a significant difference. **(C)** Bar plot for GO enrichment of DEGs. The red bars indicate GO terms for biological process. The green bars indicate GO terms for cellular component. The blue bars indicate GO terms for molecular function. Values on bars are the number of DEGs for each GO term.

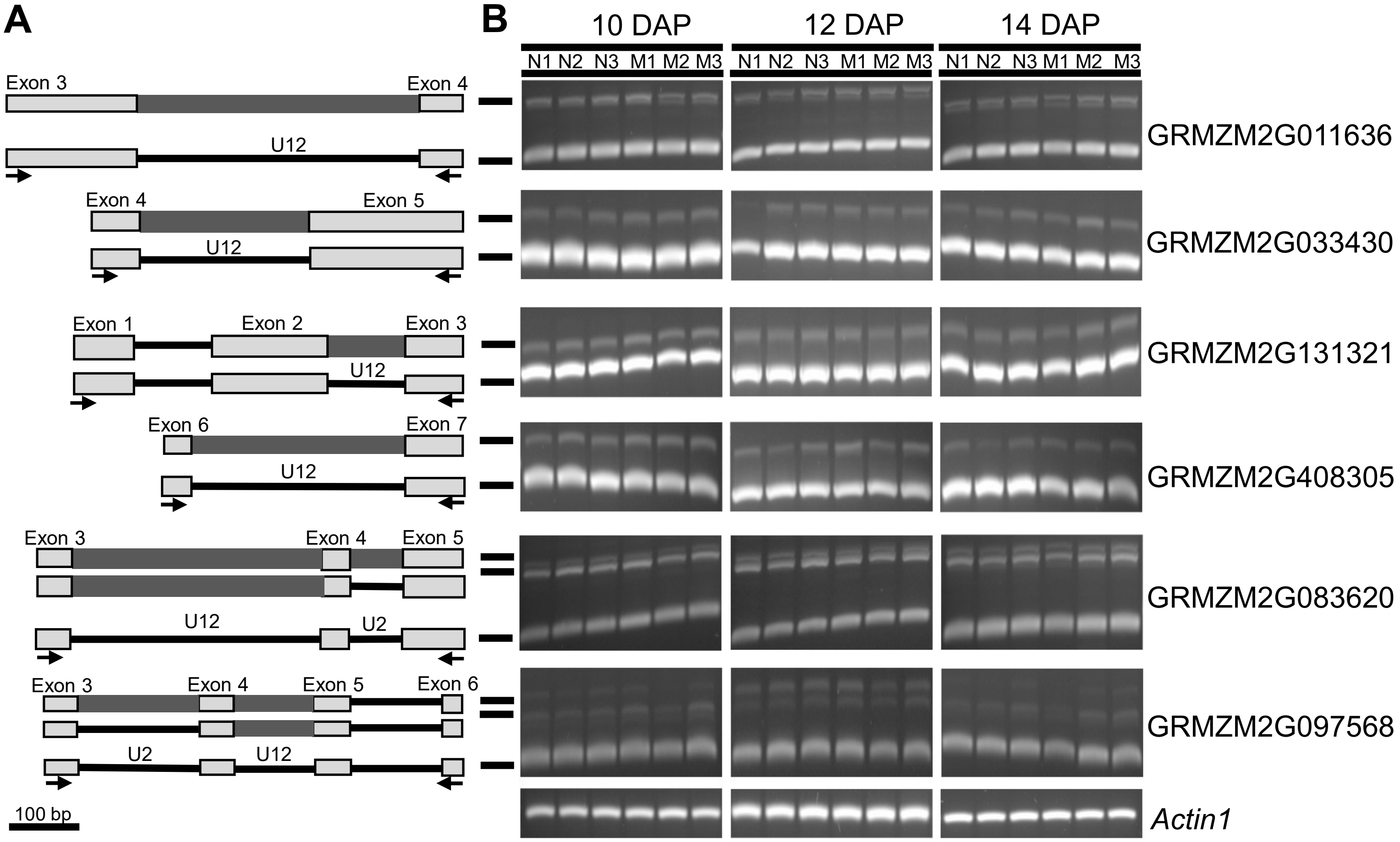


**Supplementary Figure S3. Analysis of U12-type intron splicing. (A)** Schematics show amplified products with PCR primers indicated by arrows. **(B)** RT-PCR of *rgh6* mutant (M) and normal siblings (N) RNA from endosperm tissue with three biological replicates at three developmental time points.

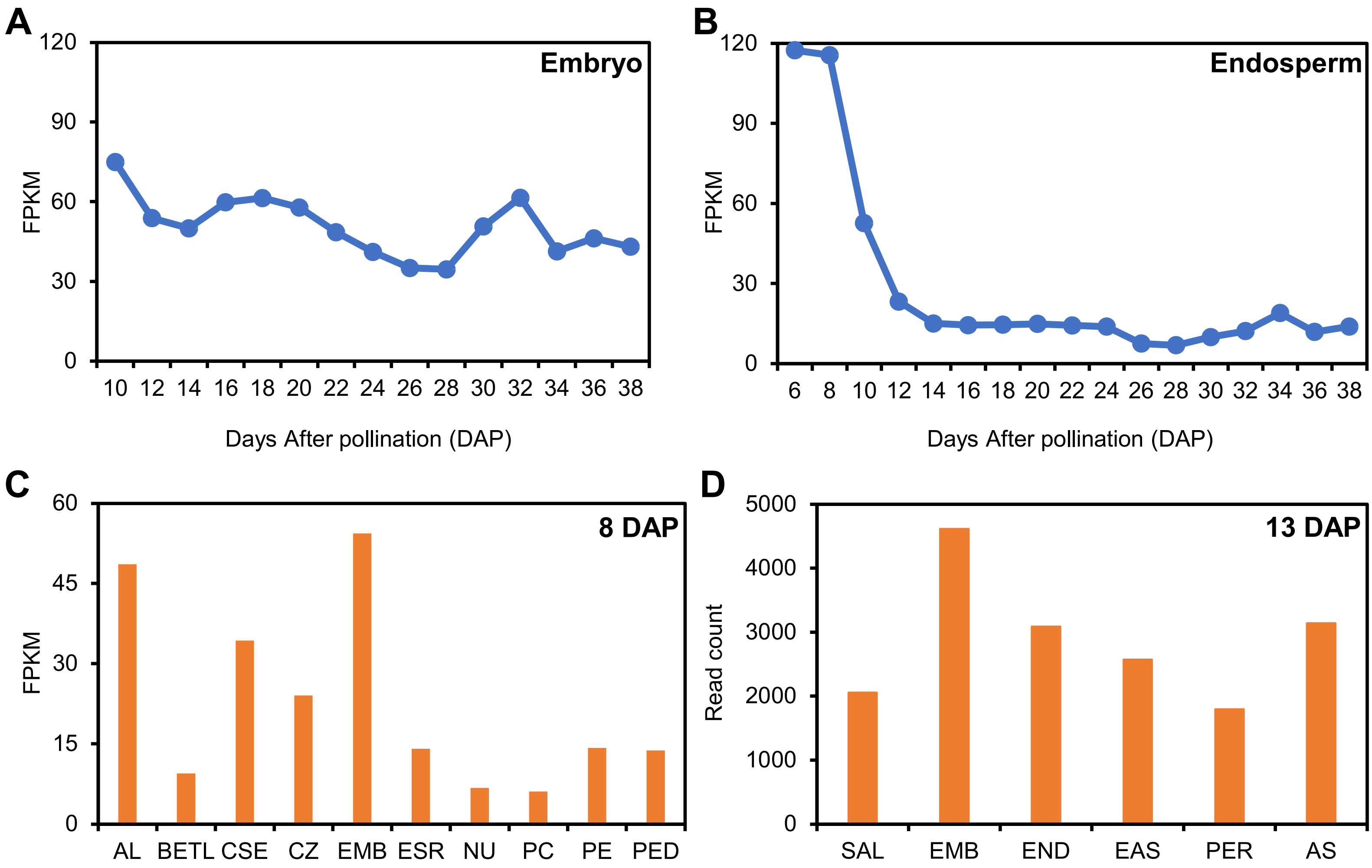


**Supplementary Figure S4. *Rgh6* transcript levels from published transcriptome studies. (A-B)** Temporal expression of *Rgh6* locus in embryo **(A)** and endosperm **(B)**. FPKM values are from Chen et al. (2014). **(C)** Spatial expression of *Rgh6* locus from dissected seed tissues at 8 DAP. FPKM values are from Zhan et al. (2015). **(D)** Spatial expression of *Rgh6* locus from dissected seed tissues at 13 DAP. FPKM values are from Doll et al. (2020).

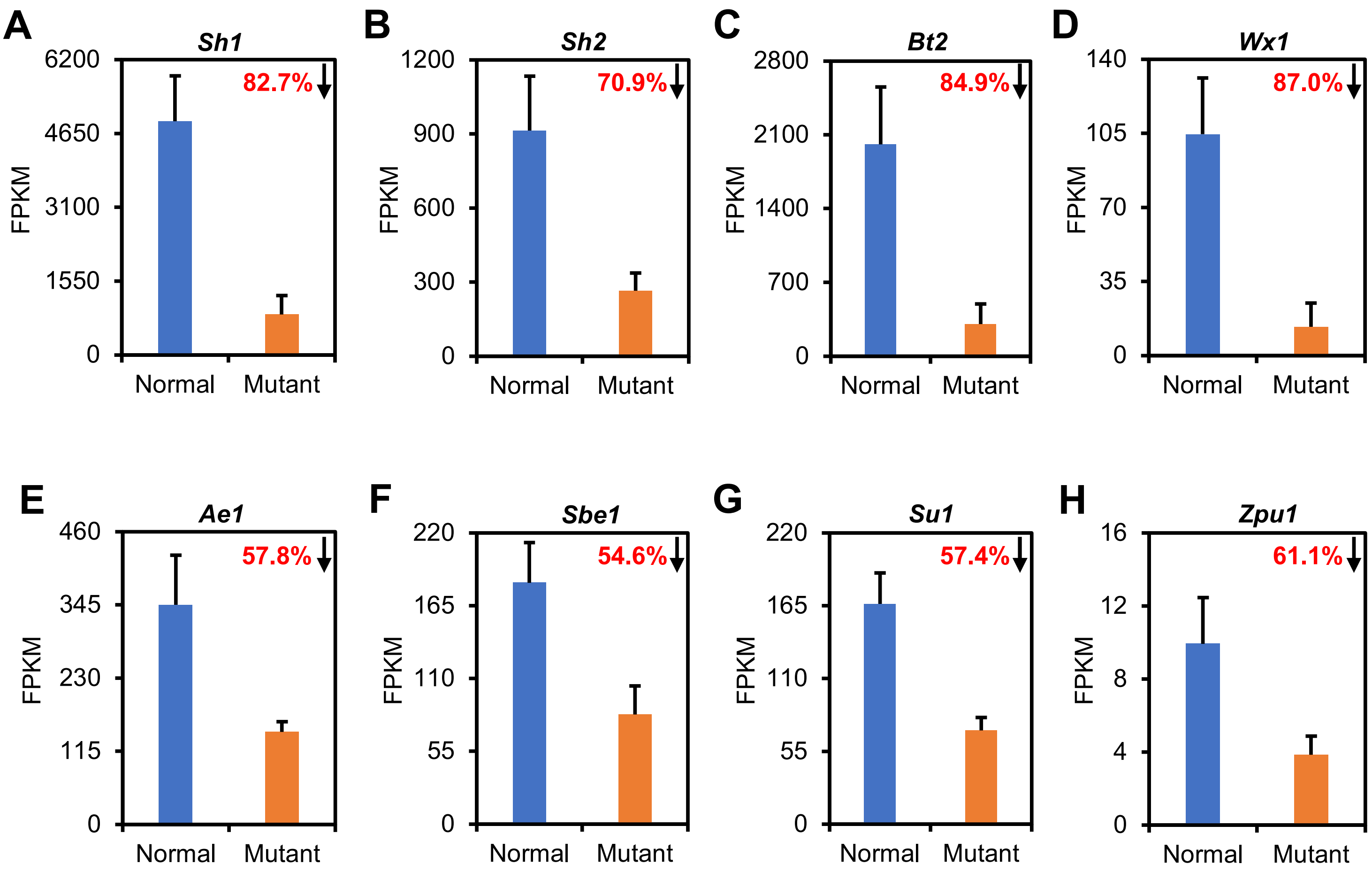


**Supplementary Figure S5. Transcript levels of starch synthesis genes in *rgh6* endosperm. (A-H)** Mean and standard deviation FPKM values are plotted from the three biological replicates of 10 DAP endosperm RNA-seq dataset for *rgh6* and normal siblings.

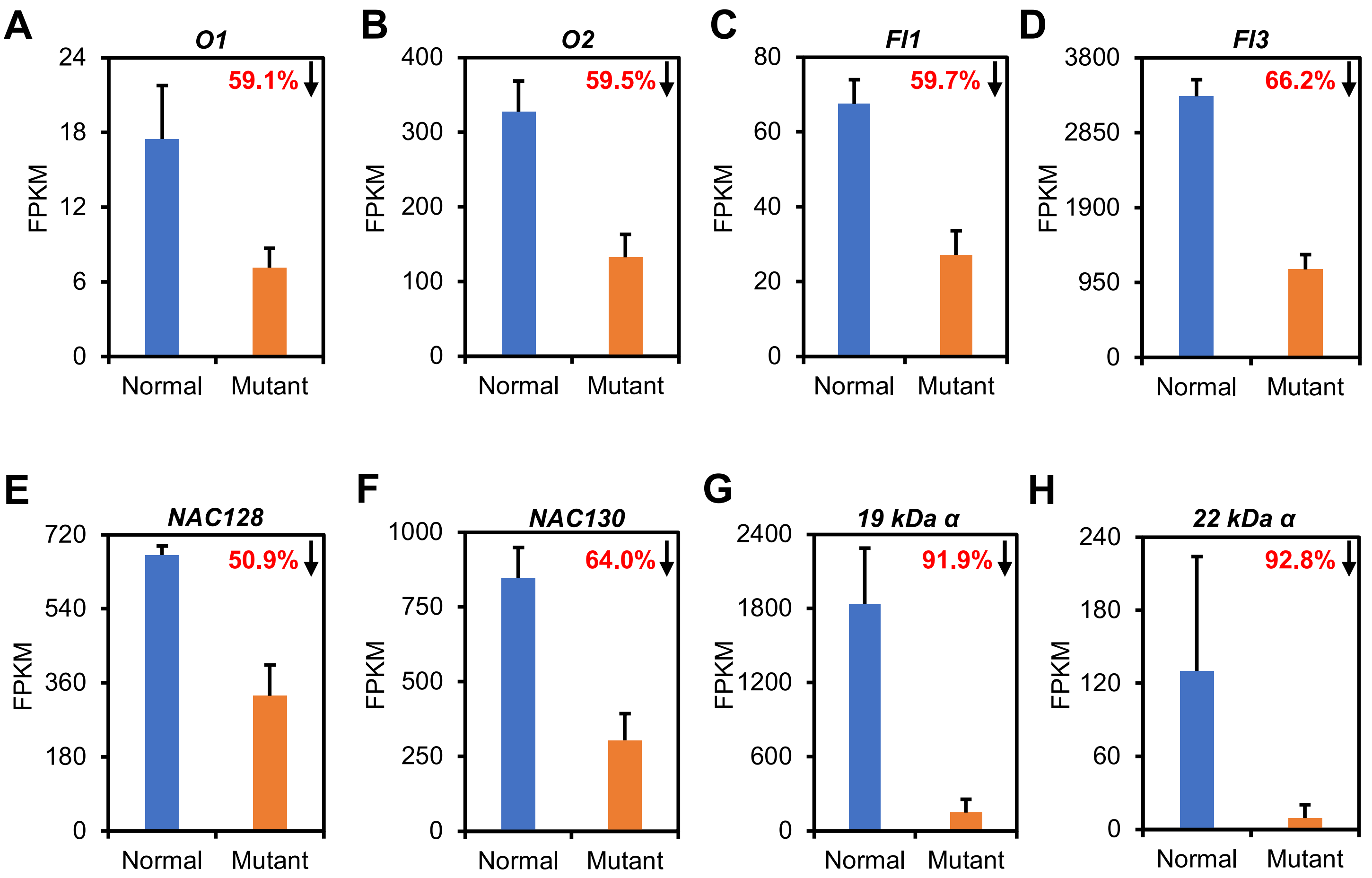


**Supplementary Figure S6. Transcript levels of zein seed storage protein related genes in *rgh6* endosperm. (A-H)** Mean and standard deviation FPKM values are plotted from the three biological replicates of 10 DAP endosperm RNA-seq dataset for *rgh6* and normal siblings.
